# The N6-methyladenosine demethylase ALKBH5 regulates the hypoxic HBV transcriptome

**DOI:** 10.1101/2023.10.31.564956

**Authors:** Senko Tsukuda, James M Harris, Andrea Magri, Peter Balfe, Peter AC Wing, Aleem Siddiqui, Jane A McKeating

## Abstract

Chronic hepatitis B is a global health problem and current treatments only suppress hepatitis B virus (HBV) infection, highlighting the need for new curative treatments. Oxygen levels influence HBV replication and we previously reported that hypoxia inducible factors (HIFs) activate the basal core promoter to transcribe pre-genomic RNA. Application of a probe-enriched long-read sequencing method to map the HBV transcriptome showed an increased abundance of all viral RNAs under low oxygen or hypoxic conditions. Importantly, the hypoxic-associated increase in HBV transcripts was dependent on N6-methyladenosine (m^6^A) modifications and an m^6^A DRACH motif in the 5’ stem loop of pre-genomic RNA defined transcript half-life under hypoxic conditions. Given the essential role of m^6^A modifications in the viral transcriptome we assessed the oxygen-dependent expression of RNA demethylases and bio-informatic analysis of published single cell RNA-seq of murine liver showed an increased expression of the RNA demethylase ALKBH5 in the peri-central low oxygen region. *In vitro* studies with a human hepatocyte derived HepG2 cell line showed increased ALKBH5 gene expression under hypoxic conditions. Silencing the demethylase reduced the levels of HBV pre-genomic RNA and host gene (CA9, NDRG1, VEGFA, BNIP3, FUT11, GAP and P4HA1) transcripts and this was mediated via reduced HIFα expression. In summary, our study highlights a previously unrecognized role for ALKBH5 in orchestrating viral and cellular transcriptional responses to low oxygen.

**Author Summary:** Oxygen levels influence HBV replication and hypoxia inducible factors (HIFs) activate HBV transcription. Long-read sequencing and mapping the HBV transcriptome showed an increased abundance of all viral RNAs under hypoxic conditions that was dependent on N6-methyladenosine modifications. Investigating the oxygen-dependent expression of RNA demethylases identified ALKBH5 as a hypoxic activated gene and silencing its expression showed a key role in regulating HBV and host gene expression under hypoxic conditions.

## Introduction

Chronic hepatitis B (CHB) is one of the world’s most economically important diseases, with 2 billion people exposed to the virus during their lifetime resulting in a global burden of >290 million chronic infections. Hepatitis B virus (HBV) replicates in the liver and chronic infection can result in progressive liver disease, cirrhosis and hepatocellular carcinoma (HCC) [1]. HBV is the prototypic member of the *hepadnaviridae* family of small enveloped hepatotropic viruses with a partial double-stranded relaxed circular DNA (rcDNA) genome. Current treatments include nucleos(t)ide analogs and interferons that suppress virus replication but are not curative largely due to the persistence of episomal HBV genomes and dysfunctional viral-specific immune responses [2].

HBV infects hepatocytes and the rcDNA genome translocates to the nucleus and is converted to covalently closed circular DNA (cccDNA) by host-DNA repair enzymes [3]. Several members of the host DNA repair pathway convert rcDNA to cccDNA that serves as the transcriptional template for all viral RNAs [4]. HBV transcribes six major RNAs of decreasing length, with a common 3’ polyadenylation signal that include: pre-core (pC) that encodes e antigen; pre-genomic (pgRNA) that is translated to yield core protein and polymerase; preS1, preS2 and S RNAs encoding the surface envelope glycoproteins and X transcript for the multi-functional x protein (HBx) [5]. pgRNA contains stem loop structures at both the 5’ and 3’ termini that bind host factors that regulate transcript stability, including zinc finger CCHC-type Containing 14 protein that recruits terminal nucleotidyltransferase 4 [6] and the zinc finger antiviral protein that regulates RNA decay [7] (reviewed in [5]). pgRNA is encapsidated and reverse-transcribed by the viral polymerase to generate rcDNA genomes that can be enveloped and secreted as infectious particles [8].

*N*6-methyladenosine (m^6^A) is the most abundant modification found on eukaryotic transcripts where it can regulate mRNA structure, stability, translation and nuclear export. m^6^A modifications are regulated by the balanced activities of m^6^A “writer” and “eraser” proteins. Adenosine is methylated by writers including two methyltransferase-like 3 (METTL3) and METTL14 [9] along with their cofactor Wilms tumor 1-associated protein (WTAP) [10]. This complex methylates adenosine residues within the consensus DRACH (D=A, G or U; R=G or A; H=A, C or U) motif which is often located near stop codons, 3’ untranslated regions and internal exons in mRNAs [11, 12]. m^6^A modifications can be removed by erasers including the demethylases, AlkB Homolog 5 (ALKBH5) and fat mass and obesity-associated protein (FTO) [13, 14]. HBV cccDNA encodes a DRACH motif present in all viral transcripts near the common 3’ polyadenylation signal in a region termed the epsilon stem loop, but notably is also found in the 5’ terminal repeat at the start of the pC and pgRNA transcripts [15]. m^6^A modified HBV RNAs are recognized by YTH domain containing protein 2 (YTHDF2) and the interferon-induced RNase, ISG20, that can process them for degradation [16]. More recently, m^6^A modified HBV RNAs were reported to be preferentially transported from the nucleus and encapsidated [17, 18]. Collectively, these studies demonstrate an important role for these post-transcriptional modifications in multiple stages of the HBV life cycle.

Oxygen concentration varies across different tissues, with the liver receiving oxygenated blood from the hepatic artery and partially oxygen-depleted blood via the hepatic portal vein, resulting in an oxygen gradient of 8-3% across the periportal and pericentral areas, respectively [19]. This oxygen gradient associates with liver zonation, a phenomenon where hepatocytes show distinct functional and structural organization across the liver [20]. Mammalian cells adapt to low oxygen through an orchestrated transcriptional response regulated by hypoxia inducible factors (HIFs): a heterodimeric transcription factor comprising alpha (HIF-1α, HIF-2α, or HIF-3α) and beta (HIF-1β) subunits [21]. Oxygen dependent prolyl-hydroxylase (PHD) enzymes hydroxylate HIF-α subunits for proteasomal degradation and hypoxic inactivation of the PHDs stabilizes HIF-α expression leading to transcription of genes involved in metabolic processes [22]. We recently showed that HIFs bind and activate HBV cccDNA transcription both in laboratory models maintained under low oxygen and in HBV transgenic mice [23]. HIFs also suppress cccDNA deamination by Apolipoprotein B mRNA Editing Catalytic Polypeptide-like 3B (APOBEC3B) [24]. To date, studies investigating the role of m^6^A modifications in the HBV life cycle have been performed under standard laboratory conditions of 18% oxygen, where HIFs are inactive. As the RNA demethylase ALKBH5 was previously reported to be regulated by HIF-1α [25], we studied the role of m^6^A modifications in the HBV life cycle under low oxygen conditions that mimic the liver. Our studies identify an essential role for m^6^A modifications in regulating the HBV transcriptome and long-read sequencing revealed oxygen- and m^6^A-dependent regulation of canonical and non-canonical viral RNAs. We also identify a role for ALKBH5 in the regulation of HIF-1α under hypoxic conditions that impacts the abundance of HBV and cellular transcripts.

## Results

### Hypoxic regulation of the HBV transcriptome is dependent on m^6^A modifications

To assess the role of m^6^A RNA modifications in HBV replication under low oxygen conditions we mutated the DRACH motifs to generate m6A-null virus as previously reported [15]. Transfecting HepG2-NTCP cells with plasmids encoding wild-type (WT) or the mutated viral genome (HBV m^6^A-null) allowed us to generate virus for infection studies. We selected 1% oxygen as this is known to stabilise HIFα subunits and activate HIF target gene transcription [23]. HBV WT or m^6^A-null infected cells were cultured at 18% or 1% oxygen for 72h and pgRNA quantified by PCR. To measure HIF-dependent regulation of pgRNAs the infected cells were treated with a prolyl-hydroxylase inhibitor (FG-4592) that activates HIF-signalling. Low oxygen or FG-4592 treatment induced a significant increase in the abundance of pgRNA in the WT infected cells, however, transcript levels were unchanged in the HBV m^6^A-null infected cells (**Fig.1A**). We noted comparable expression of the HIF-target gene carbonic anhydrase 9 (*CA9*) in WT and HBV m^6^A-null infected cells, suggesting an essential role for m^6^A modifications in the HIF induction of pgRNA.

**Figure 1.**
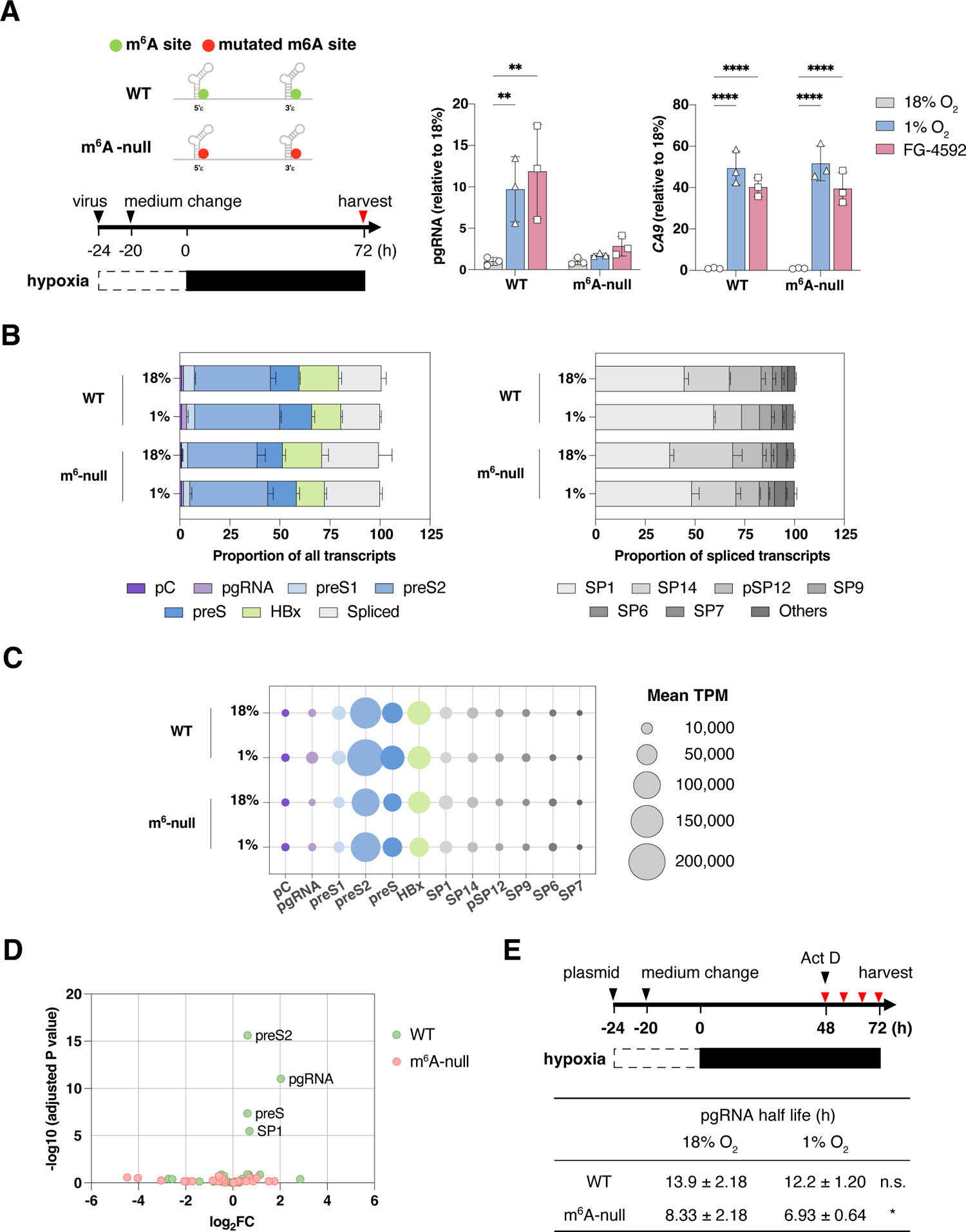
Hypoxic regulation of the HBV transcriptome is dependent on m^6^A methylation. (**A**) Schematic of HBV DRACH mutations in 5’ and 3’ pgRNA stem loops and protocol for HBV infection. HepG2-NTCP cells were infected with HBV wild type (WT) or m^6^A-null and cultured at 18%, 1% oxygen or treated with FG-4592 (30 µM) for 72h. pgRNA and HIF-regulated gene carbonic anhydrase 9 (*CA9*) transcripts were measured by qPCR. Data are expressed relative to 18% oxygen for each condition and presented as mean ± S.D. of n=3 from three independent experiments with statistical significance determined using a two-way ANOVA. * *p* < 0.05, ** *p* < 0.01, *** *p* < 0.001, **** *p* < 0.0001. (**B**) Long-read sequence analysis of HBV transcriptome. Relative frequency of canonical (left) and non-canonical (right) HBV reads in HBV WT or m^6^A-null expressing HepG2 cells cultured at 18% or 1% oxygen. (**C**) Bubble heat map denoting the HBV transcript abundance of HBV WT and m^6^A-null samples cultured at 18% and 1% oxygen. The data is presented as the mean transcripts per million (TPM) from n=3 independent samples with the exception of WT at 1% oxygen where n=2. (**D**) Differential gene expression of hypoxic regulated HBV WT and m^6^A-null RNAs, where the X axis denotes differences between samples (Log_2_-Fold Change) and the Y axis the adjusted probability of the changes seen (-log_10_ (adjusted p-value). (**E**) HBV pgRNA half-life. HepG2-NTCP cells transfected with HBV WT plasmid were incubated at 18% or 1% oxygen conditions for 48h and cells harvested at 0, 6, 12, and 24h post actinomycin D (Act D) treatment. pgRNA relative to a housekeeping gene B2M was quantified by qPCR and half-life presented as the mean ± S.D. of n=9 from three independent experiments. n.s., not significant * *p* < 0.05, by unpaired Student’s t test.

To explore the interplay between hypoxia and methylation status on the HBV transcriptome we used a probe-enrichment long-read sequencing approach [26] to map viral transcripts. We extended our earlier analytical pipeline to include all reported viral transcription start sites (TSS) [27, 28] and to map full-length transcripts. The sequenced libraries contained 47,697 - 119,071 reads and we noted a similar frequency of HBV reads among the samples, ranging from 41-55% of total reads (**Supplementary Table 1A)**. To compare the profile of HBV RNAs in the different samples we expressed the viral reads as transcripts per million (TPM) (**Supplementary Table 1B**). PreS2 RNAs were the most abundant transcript irrespective of hypoxic conditions or methylation status, with lower levels of pC, pgRNA, HBx and spliced RNAs in the HBV m^6^A-null samples compared to WT although these differences were not statistically significant (**Fig.1B-C** and **Supplementary Table 1B**). A minority of viral transcripts did not map to known TSS and were classified as unmapped, with many of these encoding an S gene open reading frame (ORF) (**Supplementary Table 1B**). Mapping the spliced transcripts identified SP1 and SP14 as the most abundant (∼15-30,000 TPM, respectively) with SP6, SP7, SP9 and pSP12 transcripts also detected (>1,000 TPM) (**Fig.1B-C, Supplementary Table 1B**). SP1 transcript levels were reduced in the HBV m^6^A-null samples compared to WT, with a concomitant increase in SP14 (**Fig.1C, Supplementary Table 1B**). Differential expression analysis identified three viral transcripts, pgRNA, preS2 and preS, together with the major spliced transcript SP1, that were significantly increased in HBV WT samples under hypoxia (adjusted p-values < 10^-6^) (**Fig.1D**). In contrast, pC, preS1 or HBx transcript levels did not increase under hypoxic conditions (**Fig.1D**). HBV m^6^A-null derived RNAs were insensitive to the low oxygen (**Fig.1D**), demonstrating a key role for m^6^A post-transcriptional modifications in defining the hypoxic HBV transcriptome.

As methylation and hypoxia can both influence RNA stability [29, 30] we measured the half-life of HBV WT or m^6^A-null derived transcripts under low-oxygen conditions by treating cultures with actinomycin D (**Fig.1E**). Given the overlapping nature of the HBV transcripts we can only accurately quantify pgRNA by qPCR. HBV m^6^A-null encoded pgRNAs had a significantly shorter half-life than WT transcripts in cells cultured under standard laboratory conditions (18% O_2_) (8.33 ± 0.71h v 13.9 ± 2.18h, *p*=0.014), consistent with reports that m^6^A modifications reduce the stability of viral RNA [15]. Hypoxia did not alter the half-life of HBV WT pgRNA but we noted a significant reduction in m^6^A-null pgRNA under these conditions (6.93 ± 0.64h v 8.33 ± 0.71h, *p*=0.044) (**Fig.1E**). In summary, these data highlight a role for oxygen-dependent processes in regulating the stability of non-methylated and methylated HBV pgRNA.

### The 5’ stem-loop DRACH motif regulates pgRNA abundance under hypoxic conditions

To investigate which m^6^A motif regulates pgRNA levels under low oxygen conditions, we transfected HepG2-NTCP cells with HBV 1.3x overlength plasmids with mutations in the DRACH motif in the 5′ loop (HBV m^6^A-5’null), the 3′ loop (HBV m^6^A-3’null) or both loops (HBV m^6^A-null) (**Fig.2A**). Transfected HepG2-NTCP cells were cultured at 18% or 1% oxygen conditions and pgRNA along with secreted HBV DNA measured (**Fig.2B**). Measuring intracellular HBV DNA at 4h post transfection demonstrated similar efficiencies of transfection (**Supplementary** Fig.1A) and we confirmed that all samples responded to the hypoxic conditions by measuring *CA9* gene expression (**Supplementary** Fig.1B). Lower levels of pgRNA were noted in the m^6^A-null and m^6^A-5’null transfected cells compared to WT or m^6^A-3’null, consistent with our earlier report showing their reduced replicative fitness (**Supplementary** Fig.1B) [15]. We observed hypoxic induction of pgRNA and extracellular HBV DNA in the WT or m^6^A-3’null transfected cells, whilst neither m^6^A-null nor m^6^A-5’null showed any oxygen-dependent modulation.

**Figure 2.**
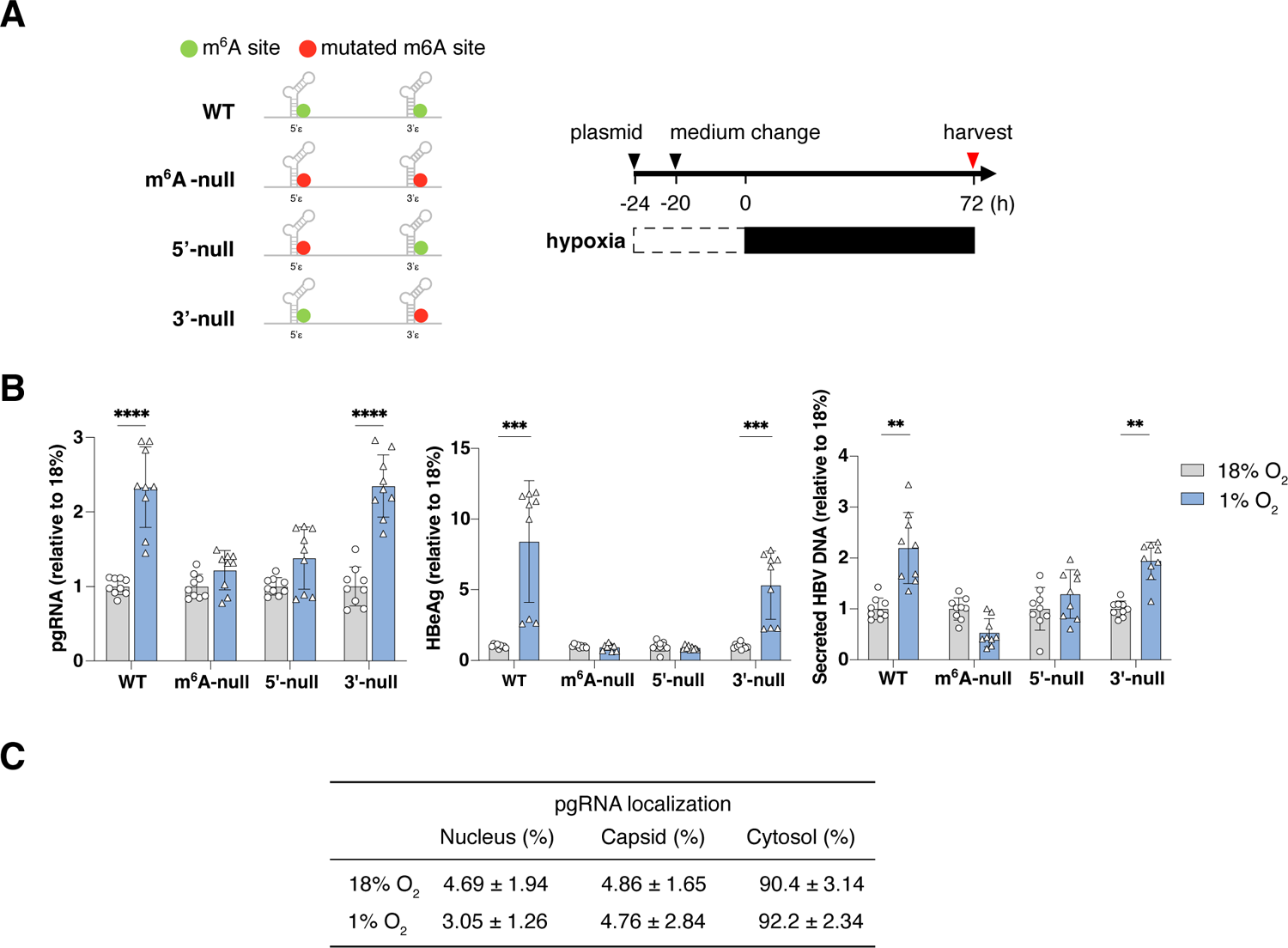
Hypoxic dependent increase in HBV pgRNA and secreted viral DNA is dependent on 5’ m^6^A methylation. (**A**) Schematic of HBV plasmids indicating the location of the A-C mutation of the 5′ and 3′ m^6^A sites in pgRNA. Circles represent WT (green) and the C mutation (red). HepG2-NTCP cells were transfected with HBV plasmids for 4h and incubated for an additional 20h at 18% oxygen, before transfer to 18% or 1% oxygen for 72h. (**B**) Intracellular pgRNA and secreted HBV DNA were quantified by qPCR and the data presented as mean ± S.D. of three independent experiments with statistical significance determined using Mann-Whitney tests, with Bonferroni correction for multiple comparisons. * *p*<0.05, ** *p* < 0.01, *** *p* < 0.001, **** *p* < 0.0001. (**C**) HepG2-NTCP cells were prepared as shown in A, and RNA extracted from the isolated cellular fractions (see methods), treated with TURBO DNase to remove any contaminating plasmid DNA and pgRNA quantified by qPCR. Data were obtained from n=5 independent samples and presented as mean ± S.D.

We were interested to understand whether the hypoxia-associated increase in extracellular HBV DNA reflected an increase in pgRNA encapsidation. HepG2 cells transfected with HBV were cultured under hypoxic conditions and the relative abundance of pgRNA in the nucleus, cytosol or within capsids measured. The majority of pgRNA was in the cytosol and this did not change under low oxygen conditions (**Fig.2C**), suggesting that hypoxia does not influence the encapsidation machinery and/or assembly processes. In summary, these data show that 5’ m^6^A modification plays an essential role in hypoxia-dependent increase in the abundance of pgRNA.

### Hypoxic activation of ALKBH5 expression impacts m6A modified HBV and host transcripts

The m^6^A demethylase ALKBH5 is a direct target of HIF-1α [25] and is hypoxic regulated in a variety of cell lines, including breast cancer and adipocyte cells [31, 32]. We investigated ALKBH5 expression in the liver using published single cell RNA-seq data of murine liver [33]. We observed an enrichment of *Alkbh5* transcripts in the low oxygen pericentral area (PC) compared to the perivenous region (PV) (**Fig.3A**). Zonal expression of the HIF target gene, N-myc downstream regulated 1 (*Ndrg1*) was noted, whereas expression of m^6^A demethylase Fat mass and obesity associated (*Fto*) gene was comparable across the liver lobule (**Fig.3A, Supplementary** Fig.2). To ascertain whether these observations translate to our *in vitro* model we cultured HepG2 cells at 18% and 1% oxygen and showed a significant increase in ALKBH5 gene expression in hypoxic cells (**Fig.3B**). To assess the functional consequences, we measured total m^6^A-RNA levels by immuno-northern blotting with an anti-m6A antibody and showed a reduction in m^6^A-modified cellular RNAs under hypoxic conditions, consistent with the increase in demethylase expression (**Fig.3C**).

**Figure 3.**
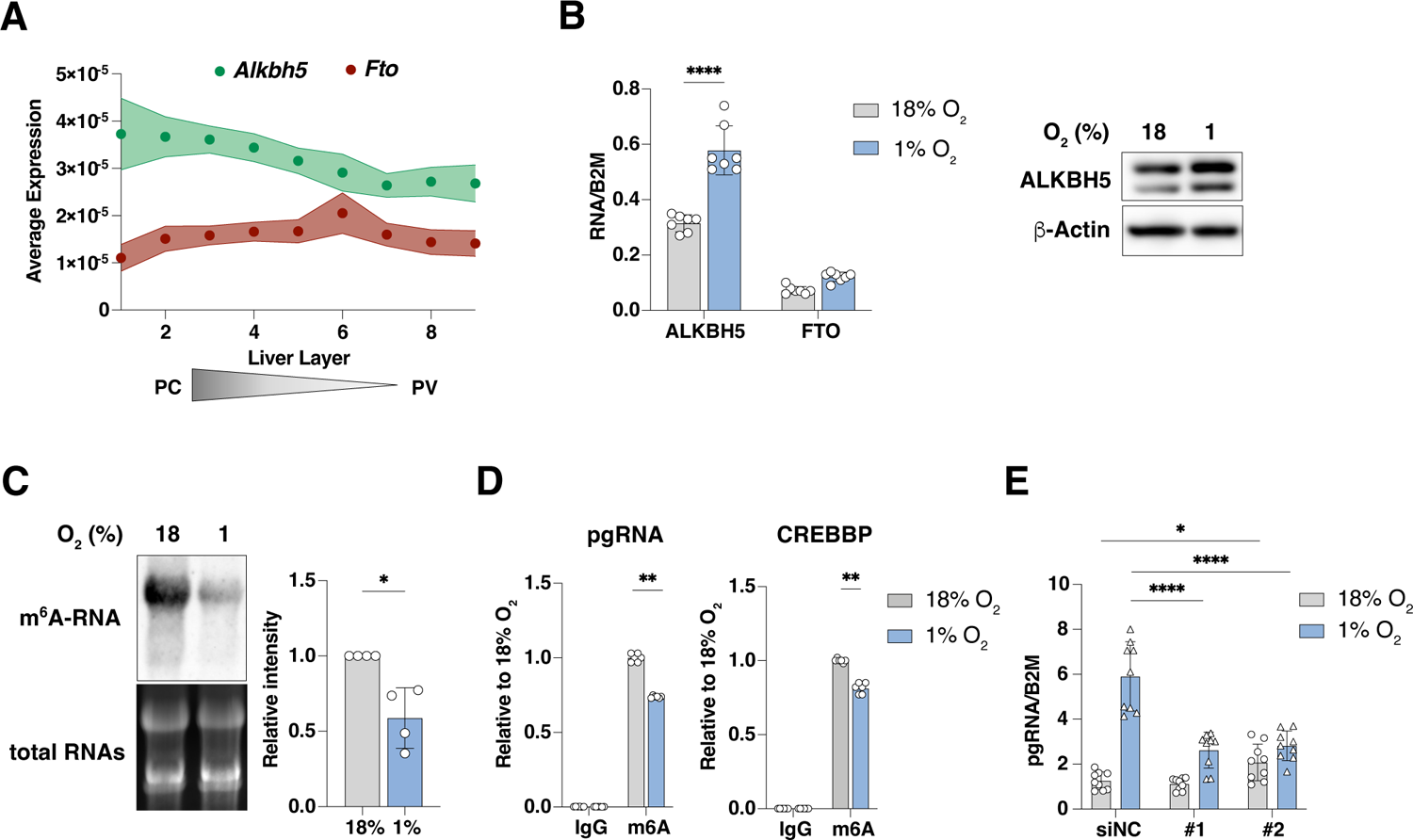
Hypoxic activation of ALKBH5 regulates methylation and abundance of HBV pgRNA. (**A**) Liver zonation of the RNA demethylases ALKBH5 and FTO based on scRNA-seq data from mouse liver [33]. (**B**) HepG2-NTCP cells were cultured at 18% or 1% oxygen conditions for 72h with ALKBH5 and FTO transcript levels (left) and ALKBH5 protein (right) measured. Data are expressed relative to a housekeeping gene *B2M* and presented as mean ± S.D. of n=4 from 2 experiments with statistical significance determined using Mann-Whitney tests, with Bonferroni correction for multiple comparisons, *** *p* < 0.01. (**C**) m^6^A-modified RNAs. RNA was extracted from HepG2-NTCP cells cultured under 18% or 1% oxygen conditions for 72h, TURBO DNase treated and subjected to immuno-northern blotting using m^6^A antibody (left). Densitometric quantification of northern blots was performed and data expressed relative to 18% oxygen. Data is shown as the mean ± S.D. of 4 independent experiments with statistical significance determined using Mann-Whitney test. * *p* < 0.05. (**D**) Quantification of methylated HBV pgRNA. HepG2-NTCP cells were transfected with HBV WT plasmid for 18h, cultured at 18% or 1% oxygen for 72h. RNA was extracted, TURBO DNase treated and subjected to Methylated RNA immunoprecipitation (MeRIP) assay using anti-m^6^A antibody. Data presented are the mean ± S.D. of n=6 samples from independent 3 experiments with statistical significance determined using Mann-Whitney tests and Bonferroni correction for multiple comparisons. ** *p* < 0.01. (**E**) HepG2-NTCP cells were transfected with siRNAs (NC; Non-targeting, ALK; ALKBH5) for 6h, followed by delivery of HBV WT plasmid and cultured at 18% or 1% oxygen for 72h. RNA was extracted, TURBO DNase digested and pgRNA quantified by qPCR. Data are expressed relative to a house keeping gene *B2M* and presented as the mean ± S.D. of n=9 samples from 3 independent experiments with statistical significance determined using Mann-Whitney tests and Bonferroni correction for multiple comparisons. * *p* < 0.05, **** *p* < 0.0001.

To investigate whether hypoxia alters HBV pgRNA methylation status we used an m^6^A–RNA immunoprecipitation (RIP) assay combined with qPCR and showed a significant reduction in precipitated pgRNA under hypoxic conditions (**Fig.3D**). As a control we PCR quantified the m6A modified CREBBP transcript and noted a reduction under hypoxic conditions. To investigate the potential role of ALKBH5 in regulating the abundance of HBV pgRNAs we silenced the demethylase with two independent siRNAs targeting different exons. We confirmed siRNA efficacy in cells cultured at 18% or 1% oxygen and siRNA #2 showed a greater reduction in ALKBH5 expression (**Supplementary** Fig.3). Silencing ALKBH5 blunted the hypoxic-associated increase in HBV pgRNA (**Fig.3E**), suggesting a role for this demethylase in regulating HBV RNAs under low oxygen conditions.

### ALKBH5 regulates HIF-**α** expression

As we previously identified a role for HIFs to bind and activate HBV transcription, we investigated the interplay between ALKBH5 and HIFs. *HIF-1α* mRNA is methylated in HepG2 cells and there was a trend albeit non-significant, for reduced amounts of anti-m^6^A precipitated transcripts under hypoxic conditions (**Supplementary** Fig.4A). HepG2 cells were transfected with siRNAs targeting ALKBH5 or an irrelevant negative control (NC) and cultured at 1% oxygen for 24h to stabilize HIF expression and activate NDRG1 expression (**Fig.4A**). Cells transfected with siALKBH5 showed a significant reduction in the expression of both HIF-α isoforms and associated NDRG1 expression, whereas *HIF-1α* transcripts were unchanged (**Fig.4A**). Of note, ALKBH5 silencing did not affect the abundance or half-life of *HIF-1α* transcripts (8.9-10.1 h) (**Fig.4B**). Silencing ALKBH5 in HepG2 cells blunted the hypoxic induction of a panel of HIF target genes (*CA9*, *NDRG1*, *VEGFA*, *BNIP3*, *FUT11*, *GP1*, and *P4HA1*), demonstrating the broad impact of this demethylase to regulate HIF-transcriptional activity (**Fig.4C**).

**Figure 4.**
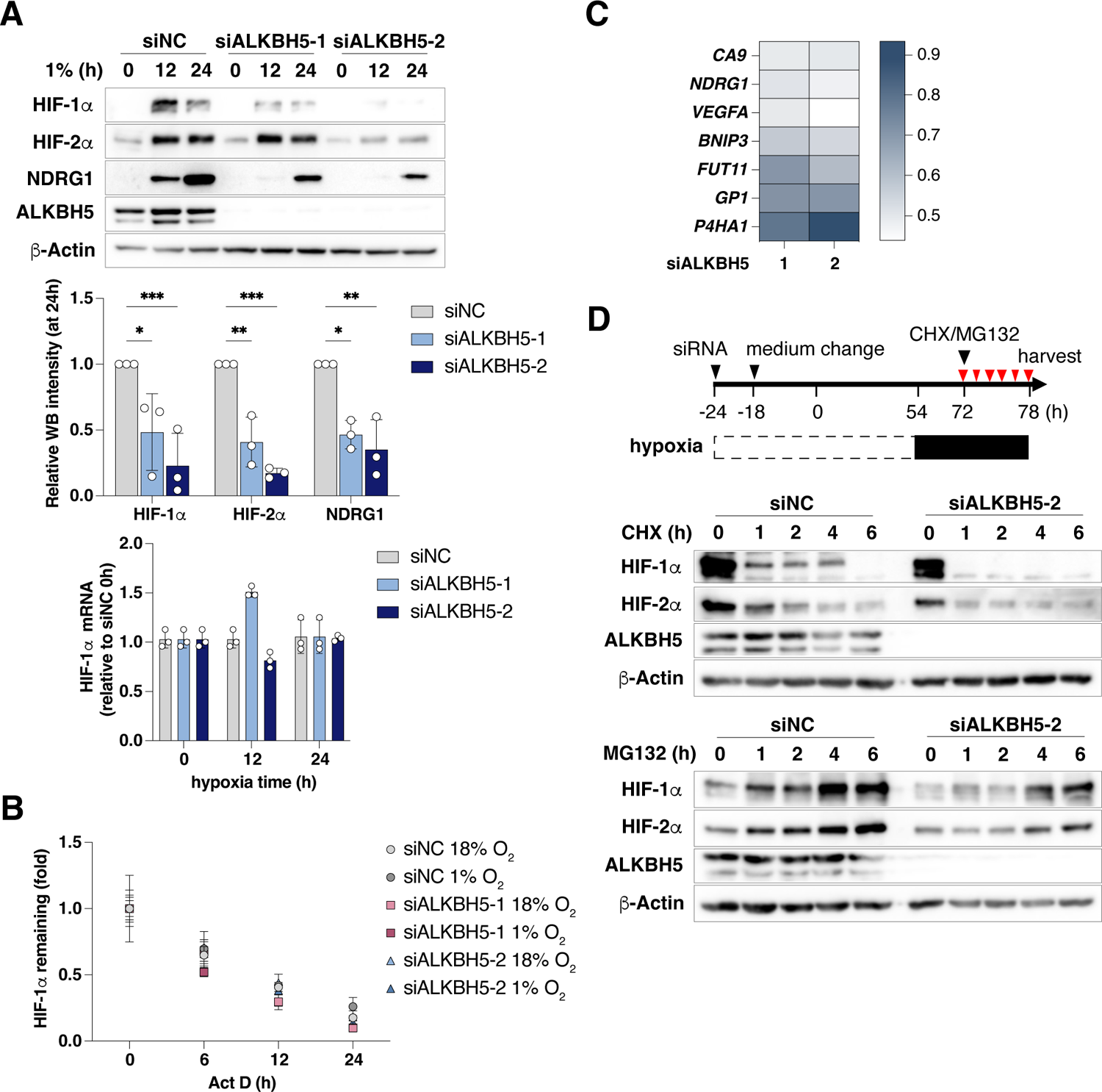
ALKBH5 regulates HIFα expression under hypoxic conditions. **(A)** HIF-α expression in ALKBH5 silenced cells. HepG2-NTCP cells were transfected with ALKBH5 specific siRNAs for 48h and cultured under 1% oxygen for 12 or 24h. Samples were probed for HIF-1α, HIF-2α, NDRG1, ALKBH5 and β-Actin by western blotting and protein expression quantified by densitometry. HIF-1α mRNA was measured by qPCR. Data is expressed relative to 24h hypoxic siNC samples and represents the mean ± S.D. from 3 independent experiments. Statistical significance was determined using a two-way ANOVA. * *p* < 0.05, ** *p* < 0.01. (**B**) HIF-1α RNA stability in ALKBH5 silenced cells. HepG2-NTCP cells transfected with ALKBH5 siRNAs were treated with Act D as shown in Fig.1C, and HIF-1α mRNA levels quantified by qPCR and expressed relative to a housekeeping gene *B2M*. Data is shown relative to 0h in each condition and expressed as mean ± S.D. from 2 independent experiments. (**C**) Expression of HIF-regulated genes in ALKBH5 silenced cells. HepG2-NTCP cells were prepared as Fig.3E, and indicated gene transcripts quantified by qPCR with data expressed relative to siNC under hypoxia and represent the mean ± S.D. of samples from 3 independent experiments. (**D**) HIF-1α protein expression in ALKBH5 silenced cells. HepG2-NTCP cells were cultured in 1% oxygen for 24h and treated with cycloheximide (CHX, 20 µg/mL) or MG132 (20 µM) for the indicated times. Samples were collected and probed for HIF-1α, HIF-2α, ALKBH5 and β-Actin by western blotting. Different exposure times for the top and bottom panels were used.

HIF-α is hydroxylated by PHDs leading to ubiquitination and targeting for proteasomal degradation. ALKBH5 silencing had no major effects on the levels of hydroxylated HIF-1α (**Supplementary** Fig.5). To assess the role for ALKBH5 in post-transcriptional regulation of HIF-α, HepG2 cells were transfected with siALKBH5 #2 as it showed the most robust silencing. The cells were cultured under hypoxic conditions for 24h before treatment with cycloheximide (CHX) to inhibit protein synthesis. CHX treatment inhibited HIF-1α and HIF-2α expression, and their degradation was accelerated in the absence of ALKBH5 (**Fig.4D**), consistent with a role for the demethylase in destabilizing both isoforms. This conclusion is further supported by treating the cells with a proteosomal inhibitor (MG132), that induced the expression of both HIF-α isoforms, and ALKBH5 silencing delaying their expression kinetics (**Fig.4D**). Collectively, these data illustrate a key role for the ALKBH5 demethylase in regulating HIF expression and function.

## Discussion

Our study shows an essential role for m^6^A RNA modifications in regulating the hypoxic HBV transcriptome. Long-read sequencing of the HBV WT infected cells allowed us to accurately map hypoxic regulated transcripts and we observed a significant increase in pgRNA, preS2, preS and SP1 transcripts, with no change in pC, preS1, HBx or other spliced mRNAs. The oxygen-dependent enrichment of viral transcripts is not dependent on their abundance and may reflect the methylation status of the different transcripts. At the present time it is not possible to profile the m^6^A methylation of viral RNAs using available long-read sequencing platforms. Given the overlapping nature of the HBV transcripts we can only reliably qPCR measure pgRNA and meRIP assays confirm that pgRNA is m^6^A modified. The observation that the hypoxia-mediated increase in HBV pgRNA is dependent on 5’-m^6^A modification suggests m^6^A specific recognition mechanisms. Wang *et al* analyzed the location of m^6^A modifications on RNAs isolated from hypoxic HeLa cells and reported that cells undergo a reprogramming of their m^6^A epitranscriptome by altering both the m^6^A level at specific sites and their distribution patterns in response to hypoxia [30]. We cannot exclude that hypoxia may alter m^6^A modified sites on HBV RNAs. Mutation of the 5’-m^6^A DRACH HBV motif at position 1907 (A to C) is unlikely to impair HIF-1α binding to hypoxic responsive elements that are located in the basal core promoter located at residues 1751-1769. RNA decay assays showed that the stability and expression of m^6^A-null pgRNA was reduced under low oxygen conditions and further experiments to image the intracellular location of methylated and non-methylated transcripts in infected cells may provide mechanistic insights.

Recent studies have highlighted an interplay between hypoxia signaling and m^6^A-post transcriptional regulatory pathways; where decreased levels of m^6^A-RNAs were seen in breast cancer cells, cardiac microvascular endothelial and cervical cancer cells [32, 34]. In contrast, increased levels of m^6^A modified RNAs were found in human umbilical vein endothelial cells, cardiomyocytes and murine hearts under hypoxic conditions that associated with increased methyltransferase METTL3 expression [35–37]. Collectively, these studies show that hypoxic modulation of m^6^A-RNA is dependent on both the tissue and cell type. Consistent with our observations, Wang *et al* reported a hypoxic reduction in m^6^A modified RNAs in human hepatoma Huh-7, HepG2 and Hep3B cell lines that was partially reversed by silencing ALKBH5 [30]. Reports that m^6^A modified RNAs are perturbed in inflammatory diseases highlight a potential therapeutic role for targeting this pathway (reviewed in [38]).

Analyzing a published single-cell RNA sequencing (scRNA-seq) data set from mouse liver [33] showed a zonation of *Alkbh5* but not *Fto*, with transcripts more abundant in the pericentral region of the liver, consistent with hypoxic regulation. ALKBH5 is a HIF target gene and its expression is induced under low oxygen conditions. ALKBH5 activity is also regulated by post-translation methylation and SUMOylation modifications that provide a further level of control over demethylase activity [39]. Our results showing reduced pgRNA levels in HBV infected ALKBH5 silenced cells under hypoxic conditions supports a positive role for this demethylase in susceptibility to viral infection. This conclusion is consistent with our earlier work that reported an increased expression of HBV antigen expressing cells in the pericentral ‘high ALKBH5’ areas of the liver in HBV transgenic mice [23, 24]. Liu *et al* reported that *Alkbh5* deficient mice were resistant to infection by a range of DNA and RNA viruses (VSV,

HSV-1 EMCV) that was mediated by an m^6^A RNA-dependent down regulation of a-ketoglutarate dehydrogenase (OGDH)-itaconate pathway that supports virus replication [40]. The authors show that Vesicular Stomatitis Virus infection targeted the ALKBH5 pathway to evade this host restriction pathway. We have limited evidence for HBV infection to alter ALKBH5 expression or activity in our experiments. Qu *et al* reported that HBx increased H3K4me3 modifications in the ALKBH5 promoter region that associated with increased demethylase expression [41]. The authors noted increased ALKBH5 expression in HBV-HCC samples, however, there were no mechanistic studies to link this directly to HBx and given the hypoxic nature of HCC (reviewed in [42]) this could align with a HIF-driven activation of ALKBH5 gene expression.

Adenosine methylation can regulate many aspects of mRNA metabolism including splicing, nuclear export, stability, and translation. Most translation events in the cell occur through recognition of the cap by the eIF4F protein complex. However, it has long been known that cellular states such as apoptosis, mitosis or the stress response can suppress cap-dependent translation allowing selected mRNAs to be translated. Recent studies reported that stress conditions, such as heat shock or amino acid starvation, promote nuclear trafficking of the m^6^A reader proteins YTHDF1 and YTHDF2 [43, 44] and increase cap-independent translation of mRNAs from m^6^A-modified 5’UTR [43, 45–47]. We found that ALKBH5 silencing reduced HIF-α expression without affecting RNA levels or half-life. Silencing of ALKBH5 did not alter the levels of hydroxylated HIF-1α, suggesting that this demethylase does not regulate PHD activity, further enzymic studies would be required to address this mechanism. There is a report showing that PBMR1, a component of the chromatin remodeler SWI/SNF, positively regulates HIF-1α translation through protein interaction between YTHDF2 under normoxic and hypoxic conditions and HIF expression was reduced when these proteins were silenced [43]. These reports are consistent with an essential role for m^6^A modified RNAs in cellular adaption to hypoxic stress. Taken together, our results support a role for ALKBH5 to regulate HIF-1α stability and transcriptional activity and in cellular adaption to hypoxic stress. Our findings on the cross-talk of ALKBH5 and HIF signalling provide mechanistic insights into cellular responses that regulate HBV replication and may be more widely applicable to other liver tropic pathogens (**Fig.5**).

**Figure 5.**
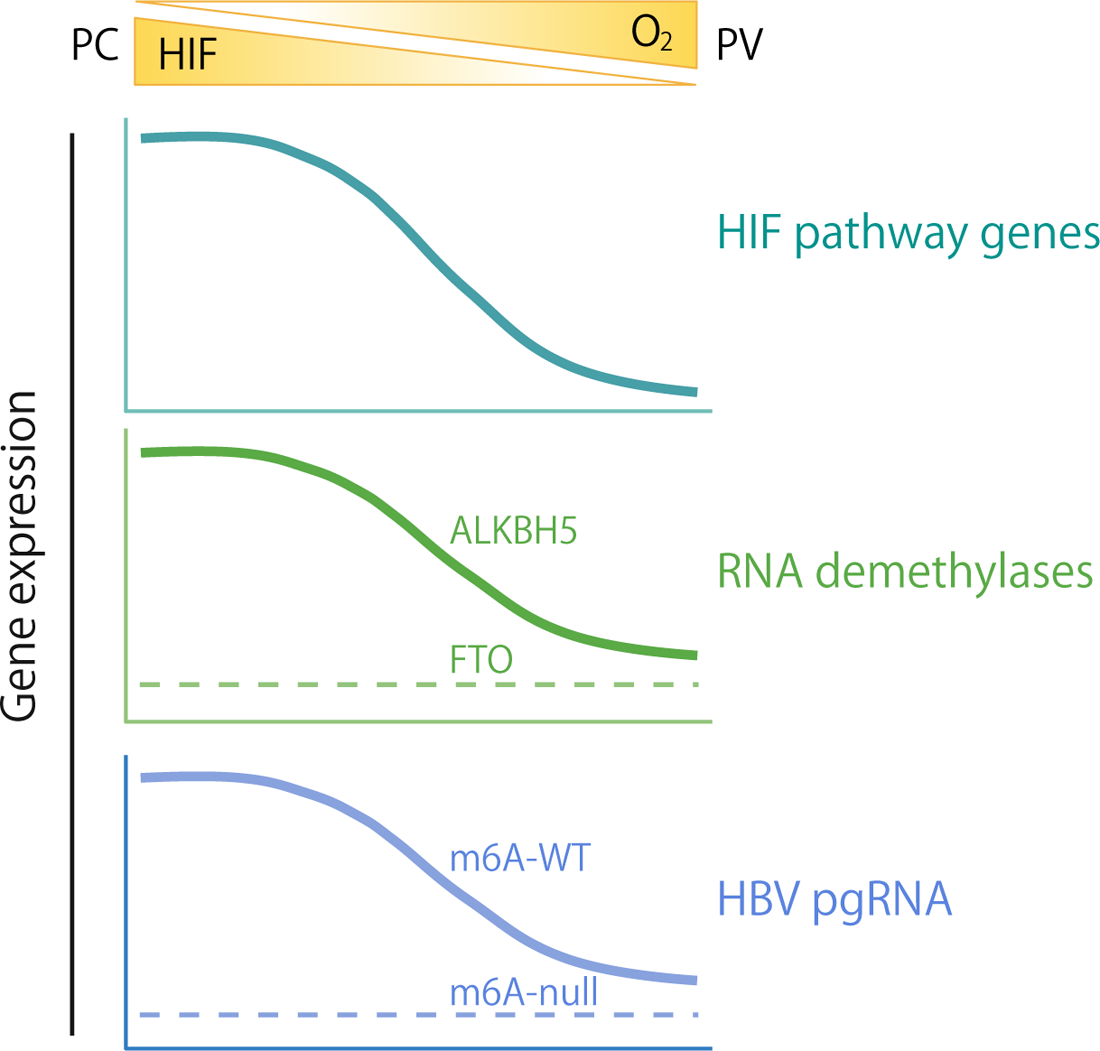
Cartoon depicting the impact of hepatic oxygen gradient on HIF signaling, RNA demethylase gene expression and m^6^A modified HBV RNAs.

## Materials and methods

### Reagents

FG-4592 was obtained from either Selleckchem or MedChemExpress. Cyclohexamide was purchased from Abcam. siRNAs were obtained from Thermo Fisher Scientific, MG132 and actinomycin D were purchased from Sigma-Aldrich. Primary antibodies used in this study are ALKBH5 (Atlas Antibodies), β Actin (Sigma), HIF-1α (BD Bioscience), HIF-2α (NOVUS), HIF-1β (NOVUS), NDRG1 (CST), anti-m^6^A polyclonal antibody (Synaptic Systems). HRP-conjugated secondary antibodies were purchased from DAKO.

### Cell lines

HepG2-NTCP cells were maintained in Dulbecco’s Modified Eagles Medium (DMEM) (ThermoFisher), containing Glutamax supplemented with 10% Fetal Bovine Serum, 50 U/ml Penicillin/Streptomycin, and non-essential amino acids. Cells were maintained in 5% CO_2_ and 18% oxygen. Hypoxic treatment of cells was carried out in a hypoxic incubator (New Brunswick Galaxy 48R, Eppendorf) or a hypoxia chamber (Invivo 400, Baker-Ruskinn Technologies) at 5% CO_2_ and 1% oxygen.

### HBV preparation and infection

HBV was prepared from the supernatant HepG2 cells transfected HBV1.3 plasmids [15] by polyethylene glycol 8000 precipitation. Briefly, culture media was mixed with 10% PEG8000/ 2.3% NaCl and incubated at 4°C for 16h, then centrifuged at 4500 rpm for 1h at 4°C. After discarding the supernatant, the pellet was resuspended in DMEM (∼200-fold concentration). The inoculum was treated with DNase at 37°C for 1h, DNA extracted and quantified by qPCR to measure HBV genome copies. HepG2-NTCP cells were seeded on collagen coated plasticware and infected with HBV (MOI of 300 or 1,000 based on genome copies) in the presence of 4% PEG8000 for 16h. Viral inoculum was removed and cells washed three times with PBS. Infected cells were maintained in 5% CO_2_ and 18% oxygen and treatments applied.

### Transfection

Plasmids were transfected into HepG2-NTCP cells using either polyethylenimine (PEI) or FuGENE HD Transfection Reagent (Promega) according to the manufacturer’s protocol. siRNAs for ALKBH5 (1; s29686 and #2; s29688) and non-targeting control (4390844) were obtained from Thermo Fisher Scientific and transfected using DharmaFECT 4 Transfection Reagent (Horizon Discovery) according to the manufacturer’s protocol.

### Quantitative PCR

Total cellular RNA was extracted using an RNeasy kit (Qiagen), then treated with TURBO DNA-free (Thermo Fisher Scientific), and the RNA reverse transcribed using a cDNA synthesis kit (PCR Biosystems) according to the manufacturer’s protocol. Cellular DNA was extracted using QIAamp DNA kit (Qiagen). Gene expression was quantified using a SyGreen Blue Mix (PCR Biosystems) with the oligonucleotides listed below and a qPCR program of 95°C for 2 min followed by 45 cycles at 95°C for 5 sec, 60°C for 30 sec. Changes in gene expression were calculated by the ΔΔCt method relative to a housekeeper gene, β2-microglobulin (B2M).

### PacBio long read sequencing and analysis

HBV specific oligonucleotide enrichment and subsequent long-read sequencing and analysis was performed as reported [48]. Briefly, RNAs were prepared from normoxic or hypoxic HepG2-NTCP cells transfected with HBV HBV1.3 WT or m6-null constructs and 150ng of RNA reverse transcribed with barcoded sequencing primers. Oligonucleotide enrichment of HBV RNAs was performed as previously described [49]. Samples were sequenced using a Sequel II instrument to generate a PacBio ‘Hifi library’. Reads were mapped to the HBV reference genome (genotype D3, ayw strain) using minimap2 [50, 51]. HBV reads were assigned to previously reported transcription start sites to identify canonical transcripts [27] and splice junctions enumerated to identify non-canonical RNAs [51], incomplete sequences that did not encode the expected length of transcript were discounted. Differential gene expression was performed using the Voom function in Limma (Bioconductor EdgeR script). The sequencing data is available via the SRA at NCBI (BioProject ID: PRJNA1000182).

### Fractionation of HBV pgRNA in the nucleus, cytosol and core particles

HepG2-NTCP cells transfected with HBV1.3 plasmid were washed with PBS and lysed in 50 mM Tris-HCl, pH 8.0, and 1% NP-40 with protease inhibitor cocktail. After incubating cells at 4°C for 20 min in the culture plate, the lysate was centrifuged for 5 min at 14,000 rpm. The pellet was the nuclear fraction and RNA extracted using Trizol. HBV core particles were isolated from the supernatant according to the protocol described by Belloni et al [52]. Briefly, 100 mM CaCl2, DNase I, and RNase A were added to the supernatant and incubated for at 37°C for 2 h. The supernatant was the incubated in 5 mM EDTA, 7% PEG8000,1.75M NaCl, at 4°C for 2 h. After centrifugation at 13,000 rpm for 30 min at 4°C, the supernatant was discarded and the capsid-containing pellet resuspended in TNE buffer (10 mM Tris-HCl (pH 8) 1mM EDTA, proteinase K), and RNA was extracted using Trizol. To calculate pgRNA levels in the cytosol, total HBV RNAs were extracted from the cells using Trizol and pgRNA quantities in the nucleus and capsid subtracted after quantifying by qPCR.

### Extracellular HBV DNA quantification

Extracellular HBV DNA was quantified according to the protocol described previously [53]. Briefly, culture supernatant was treated with DNase I (Thermo Fisher Scientific) at 37°C for 60 minutes, then treated with 2x lysis buffer (100 mM Tris-HCL (pH7.4), 50 mM KCl, 0.25% Triton X-100, and 40% glycerol) containing 1mM EDTA. HBV DNA was amplified by qPCR using primers for HBV rcDNA and quantified against a DNA referent standard curve.

### SDS-PAGE and western blot

Samples were lysed in RIPA buffer (50 mM Tris (pH 8.0), 150 mM NaCl, 1% Nonidet P-40, 0.5% sodium deoxycholate, and 0.1% sodium dodecyl sulphate) supplemented with protease inhibitor cocktail tablets (Roche). 4x Laemmli reducing buffer was added to samples before heating at 95°C for 10 min. Proteins were separated on 8 or 14% polyacrylamide gel and transferred to activated 0.45 µm PVDF membranes (Amersham, UK). Membranes were blocked in 5% skimmed milk and proteins detected using specific primary and HRP-secondary antibodies. Proteins were detected using SuperSignal West Pico chemiluminescent substrate kit (Pierce) and images collected on a G:Box mini (Syngene).

### Immuno northern assay

10 µg of RNA was electrophoresed in a 1 % MOPS agarose gel containing 2.2 M formaldehyde. 18 S and 28 S ribosomal RNA species were visualized under UV light after electrophoresis to verify the amount of RNA loaded and to assess degradation. After denaturation with 50 mM NaOH for 5 min, RNAs were transferred to a nylon membrane by capillary transfer using 20× SSC buffer. After UV crosslinking, the membrane was blocked in 5% skimmed milk and incubated with an anti-m^6^A polyclonal antibody (Synaptic Systems) and HRP-secondary antibodies. Signals were realised using a SuperSignal West Pico chemiluminescent substrate kit (Pierce) and images collected on a G:Box mini (Syngene).

### MeRIP

Total cellular RNA was extracted using an RNeasy (Qiagen) kit and a TURBO DNA-free Kit (Thermo Fisher Scientific), the RNA was then further purified using RNeasy kit. 2 µg of RNA was incubated overnight at 4°C with protein A agarose beads treated with Rabbit IgG or anti-m^6^A polyclonal antibody (Synaptic Systems) in MeRIP buffer (Tris-HCl (pH 8.0),150 mM NaCl,0.1% NP-40, 1mM EDTA) supplemented with RNase inhibitor (Promega). Beads were washed 5 times with MeRIP buffer and bound RNA eluted by 6.7 mM m^6^A sodium salt. Eluted RNA was purified using a Qiagen RNA extraction kit and quantified by qPCR. Quantities of HBV RNA, *CREBBP*, *HPRT1*, and *HIF-1α* were calculated relative to input total RNA.

## Abbreviations

HBV: hepatitis B virus

HCC: hepatocellular carcinoma

rcDNA: relaxed circular DNA

cccDNA: covalently closed circular DNA

pgRNA: pregenomic RNA

HBe: hepatitis B e antigen

HBx: HBV X protein

m^6^A: N6-methyladenine

HIF: hypoxia-inducible factor

NTCP: sodium taurocholate cotransporting polypeptide

2OG: 2-oxoglutarate

MeRIP: methylated RNA immunoprecipitation

CREBBP: CREB binding protein

CA9: carbonic anhydrase 9

NDRG1: N-myc downstream-regulated gene 1

PHD: prolyl hydroxylase domain proteins

VHL: von Hippel-Lindau disease tumor suppressor

METTL3/14: methyltransferase-like protein 3/14

ALKBH5: α-ketoglutarate-dependent dioxygenase alk B homolog 5

FTO: fat mass and obesity-associated protein

YTHDF1-3: YTH domain containing family 1-3

YTFDC1-2: YTF domain containing1-2

WB: western blotting

siRNA: small interfering RNA

PCR: polymerase chain reaction

qPCR: quantitative PCR.

## Acknowledgements

We thank Ulla Protzer (TUM, Germany) for providing HBV stocks, Stephan Urban (University of Heidelberg) for HepG2-NTCP cells, Azim Ansari for the enrichment probes and Esther Ng for advice in analysing and mapping HBV sequences. The McKeating laboratory is funded by a Wellcome Investigator Award 200838/Z/16/Z, Chinese Academy of Medical Sciences Innovation Fund for Medical Science, China (grant number: 2018-I2M-2-002), AM is supported by the John Black Foundation. AS is supported by and NIH AI139234 grant.

## Supporting information

**Supplementary Figure 1. qPCR measurement of intracellular HBV DNA and pgRNA in transfected cells.** HepG2-NTCP cells were transfected with HBV plasmids (4h) and cultured at 18% or 1% O_2_ for 72h. (**A**) Total cellular DNA was extracted at 4h post-transfection and HBV DNA measured, normalized to *PrP* housekeeping gene and expressed relative to HBV WT. Data are presented as mean ± S.D. of n=3 from one of independent 3 experiments. (**B**) HBV pgRNA and *CA9* mRNAs were quantified by qPCR, expressed relative to *B2M* housekeeping gene with the data presented as mean ± S.D. of n=9 samples from 3 independent experiments, with statistical significance assessed using a non-parametric (Kruskall-Wallis) ANOVA, * *p* < 0.05, ** *p* < 0.01 *** *p* < 0.001, **** *p* < 0.0001.

**Supplementary Figure 2. Liver zonation profile of a HIF pathway gene.** Liver zonation profile of a HIF pathway gene, *Ndrg1*, based on scRNA-seq data of mouse liver [33].

**Supplementary Figure 3. Efficiency of ALKBH5 siRNAs**. Efficiency of siRNA #1 and #2 targeting ALKBH5. HepG2-NTCP cells transfected with siRNA (NC; Non-targeting, ALK; ALKBH5) for 6h and cultured at 18% or 1% O_2_ for 72h and RNA extracted. *ALKBH5* RNA and protein levels were detected by qPCR and western blotting, respectively. Statistical significance was assessed using Mann-Whitney tests, with Bonferroni correction for multiple comparisons, *** *p* < 0.001.

**Supplementary Figure 4. *HIF-1*α mRNA methylation.** Methylated RNA immunoprecipitation (MeRIP) assay. HepG2-NTCP cells were incubated at 18% or 1% oxygen for 72h. RNA was extracted, treated with TURBO DNase and incubated overnight at 4 °C with protein A agarose beads with IgG or m^6^A antibody. The beads were washed 5 times, and bound RNA eluted in 6.7 mM m^6^A sodium salt, purified using a Qiagen RNA extract kit and *HIF-1*α transcripts measured by qPCR. Data are expressed relative to input total RNA and represent the mean ± S.D. of n=4 samples from 2 independent experiments.

**Supplementary Figure 5. Hydroxylated HIF-1α protein expression in ALKBH5 silenced cells.** HepG2-NTCP cells were cultured at 1% oxygen for 24h and treated with MG132 (20 µM) for 4h. Samples were collected and probed for HIF-1α, ALKBH5 and β-Actin by western blotting.

**Supplementary Table 1**. HBV transcriptome analysis by long-read sequencing. (A) HBV reads between samples. (B) HBV transcript frequencies.

