## Supplementary Figures. for "The N6-methyladenosine demethylase ALKBH5 regulates the hypoxic HBV transcriptome"

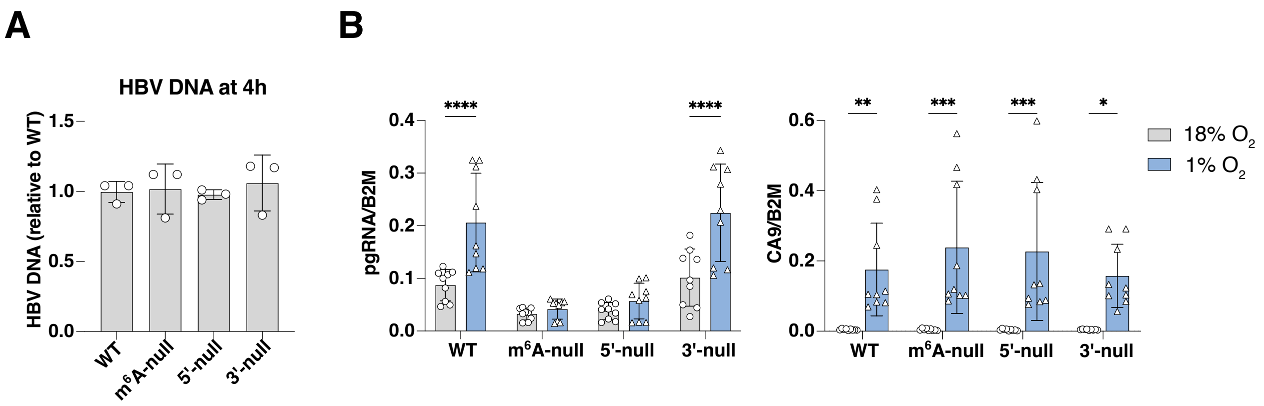


### Supplementary Figure. 1

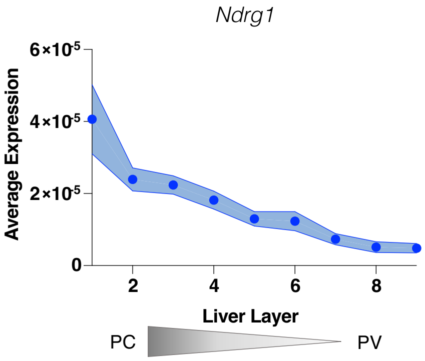


### Supplementary Figure. 2

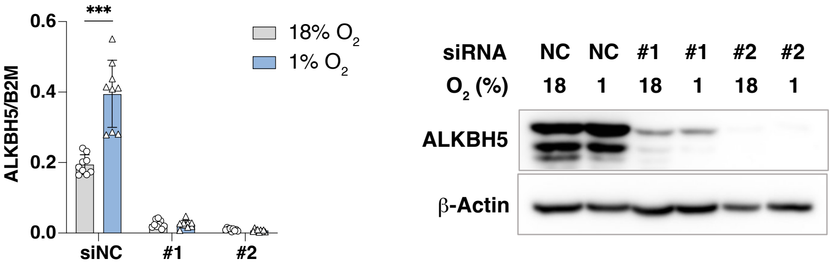


### Supplementary Figure. 3

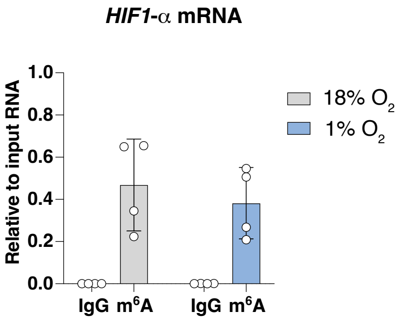


### Supplementary Figure. 4

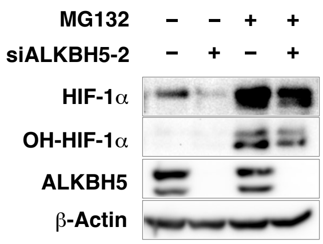


### Supplementary Figure. 5
